# Open-top axially swept light sheet microscopy

**DOI:** 10.1101/2020.12.27.424265

**Authors:** Bumju Kim, Myeongsu Na, Soohyun Park, Kitae Kim, Jeong-Hoon Park, Euiheon Chung, Sunghoe Chang, Ki Hean Kim

## Abstract

Open-top light sheet microscopy (OT-LSM) is a specialized microscopy for the high throughput cellular imaging of large tissues including optically cleared tissues by having all the optical setup below the sample stage. Current OT-LSM systems had relatively low axial resolutions by using weakly focused light sheets to cover the imaging field of view (FOV). In this report, open-top axially swept LSM (OTAS-LSM) was developed for high-throughput cellular imaging with the improved axial resolution. OTAS-LSM swept the tightly focused excitation light sheet across the imaging FOV by using an electro tunable lens (ETL) and collected emission light at the focus of light sheet with a camera in the rolling shutter mode. OTAS-LSM was developed by using air objective lenses and a water prism for simplicity and it had on-axis optical aberration associated with the mismatch of refractive indices between air and immersion medium. The effects of optical aberration were analyzed by both simulation and experiment. The image resolutions were 1.5-1.6μm, and approximately 140% of the aberration-free theoretical values. The newly developed OTAS-LSM was applied to the imaging of optically cleared mouse brain and small intestine and it could visualize neuronal networks in the single cell level. OTAS-LSM might be useful for the high-throughput cellular examination of optically cleared large tissues.

## 1. Introduction

High-speed optical microscopy methods, which can do the cellular imaging of large tissue specimens, are useful for both the biological study of neuronal network and the cellular examination of tissue specimens together with optical clearing. There are various optical microscopy techniques which can do the large tissue imaging. Light sheet microscopy (LSM) was originally developed for the high-speed 3D imaging of tiny model organisms in developmental biology with minimal photo-damage by introducing excitation light as a sheet from the side and conducting planar imaging [1–3]. LSM was later applied to the imaging of the larger tissues such as optically cleared mouse brain [4]. However, LSM showed limited performance with large samples due to the degradation of image resolution with the increase of sample size. To overcome the limitation of LSM, axially swept LSM (AS-LSM) methods were developed [5–9]. AS-LSM used a tightly focused light sheet and axially swept the light sheet across the imaging FOV. Emission light generated at the focus was collected by an imaging camera in the rolling shutter mode. AS-LSM methods have been applied to the imaging of large samples such as mouse brain [7, 9], bone marrow [7], spinal cord [8] and chicken embryo [9]. However, these LSM methods still have the limitations of sample size by keeping the configuration of side illumination and top or bottom side imaging. There have been various LSM methods developed in different optical configurations for the large sample imaging [10-12]. LS theta microscopy had both the illumination and imaging arms on one side so that large tissue samples could be placed on the other side and imaged without size limitation [10]. LS theta microscopy had the illumination arm at an oblique theta angle (less than 90 degree) and the imaging arm normal with respect to the sample surface, and the light sheet scanned in the imaging plane. LS theta microscopy had relative high image resolutions in the lateral direction and could use the full working distance of the objective lens. However, the current LS theta microscopy was in the upright configuration and had limitations in handling and imaging the tissue samples. Swept confocally-aligned planar excitation (SCAPE) microscopy was another LS microscopy without sample size limitation [11, 12]. SCAPE microscopy was a single objective lens system which had both the oblique light sheet illumination and emission light collection along the light sheet implemented. The tilted image plane by the light sheet was rotated by using an additional relay imaging arm in the matched tilted angle. Although SCAPE microscopy was simple by using a single objective lens, the image resolution varied depending on the direction and spatial location and it could be sensitive to optical aberration. Open-top LSM (OT-LSM) had separate illumination and imaging arms in the inverted configuration [13-20]. The two arms were oriented at 45° with respect to the sample surface: the illumination arm at +45□° and the imaging arm at −45□°. OT-LSM could do the planar imaging continuously with sample translation and was good for high throughput imaging. The oblique illumination and imaging with respect to the sample surface in the OT-LSM required interfacing devices such as the water prism [13, 14], solid immersion lens (SIL) [15–18] and solid immersion meniscus lens (SIMlens) [19, 20] for the normal incidence of illumination and emission light to the medium and the minimization of optical aberration associated with the oblique incidence of light on the interface. Standard aberration correction based on the wavefront sensing and correction, and the use of immersion objective lens without the interfacing devices were implemented for high-resolution imaging. Although the lateral resolution was improved by various methods, the axial resolution remained the same by using the weakly focused light sheet (> 4μm in width) covering the entire FOV in focus. The axial sweeping method could be adapted to improve the axial resolution as well.

In this report, we developed open-top axially swept light sheet microscopy (OTAS-LSM) for the high-throughput cellular imaging of optically cleared tissues with the improved axial resolution. OTAS-LSM used an electro-tunable lens (ETL) to sweep the excitation light sheet across the FOV and collected emission light from the sample by using a camera in the rolling shutter mode. The current OTAS-LSM was developed using air objective lenses in both the illumination and imaging arms and a water prism for the normal incidence of light onto the immersion medium for simplicity so that it had inherent on-axis optical aberration. The effect of aberration on image resolution was analyzed by both simulation and experiment. After the characterization, OTAS-LSM was applied to the optically cleared mouse brain and small intestine specimens for testing its ability of high-throughput cellular imaging.

## 2. Method

### 2.1 System configuration of OTAS-LSM

OTAS-LSM was based on the configuration of water-prism OT-LSM. An ETL (EL-16-40-TC-VIS-5D-M27, Optotunes) was included in the illumination arm for the axial sweeping of excitation light sheet, and emission light from the sample was collected at a sCMOS camera (ORCA-Flash4.0 V3 Digital CMOS camera, Hamamatsu) in the rolling shutter mode. OTAS-LSM used 10x air objective lenses (MY10X-803, NA 0.28, Mitutoyo) in both the illumination and imaging arms, which were located below the sample holder and oriented at +45° and −45° with respect to the sample surface. OTAS-LSM had the custom water prism filled with refractive index (RI) matching solution (C match, 1.46 RI, Crayon Technologies, Korea) for the normal light incidence onto the clearing solution.

Various schematics of OTAS-LSM system are shown in Fig. 1. A simplified optical configuration of the system, the ray tracing of the excitation light sheet in the illumination arm in two different cross sections of x-z and y-z planes, detail schematics of the water prism and specimen are shown. Excitation light was from either 488nm or 532nm CW lasers (Sapphire 488 LP −100, Coherent; LSR532NL-PS-II, JOLOOYO) and was delivered to the system via fiber coupling (SM1FC, Thorlabs; P1-460Y-FC-1, Thorlabs; HPUCO-23-532-sm-4.5as, Oz Optics). Excitation light from the fiber was collimated by a lens (L1, AC254-040-A, Thorlabs). The collimated excitation beam was converted to the sheet beam by a cylindrical lens (LJ1695RM-A, Thorlabs). The sheet beam was relayed to the sample by using two lens pairs of L2 (AC254-100-A, Thorlabs) and L3 (AC254-200-A, Thorlabs), and L4 (AC254-150-A, Thorlabs) and the objective lens. The ETL was place in between the first lens pair (L2 and L3) to control the collimation of sheet beam in the Fourier plane and to change the axial position of the sheet beam in the sample. The width of excitation sheet beam before the objective lens was approximately the same size as the back aperture of the objective lens to generate light sheet beam by using the full numerical aperture (NA) of the objective lens (0.28NA). The light sheet after the objective lens was designed to be approximately 2.13mm wide and 1.0μm thick at the beam waist in full width at half maximum (FWHM), and the focus of light sheet was 25 μm long. Excitation light from the air objective lens entered the sample through the water prism. The water prism formed the interface of RI matching solution for the normal incidence of excitation light into the medium. Excitation light passed through the RI matching solution in the water prism, a quartz coverslip (Round quartz coverslip, Ted Pella) at the bottom of the sample holder, and then got focused in the sample immersed in the RI matching solution. The quartz coverslip was used owing to its matched RI (1.46) with the one of RI matching solution. Emission light generated in the sample by the excitation light sheet was collected at the imaging arm, oriented orthogonally with respect to the illumination arm. Emission light in the sample passed through the sample holder and the water prism and was collected by the other air objective lens. After the objective lens, emission light passed through an emission filter (ET525LP and ET542LP, Chroma) and a camera lens (A17 AF 70-300mm F4-5.6 Di LD MACRO 1:2, Tamron) and then was collected at the sCMOS camera. The camera lens worked as a variable tube lens with the adjustable focal length from 180mm to 300mm. The sCMOS camera collected emission light in the rolling shutter mode. The speed of the rolling shutter and the exposure time of the camera were adjusted to be matched with the sweeping speed of excitation light sheet in the sample for confocal gating. The collected image data was transferred to a data acquisition computer via a frame grabber (Firebird camera link frame grabber, Active silicon). The translation stage (MS-2000 FLAT-TOP XY automated stage, ASI) translated the specimen between each frame. The planar imaging of the specimen at the oblique angle with respect to the sample surface was conducted continuously with the linear sample translation.

The custom water prism was used in the OTAS-LSM. The water prism was in the shape of a right-angled prism made of aluminum and regular coverslips were glued onto the right-angled sides as the windows. Its size was 8.5 mm long in the right-angled side. The water prism held the RI matching solution. The imaging FOV was approximately 1mm x 1mm with 2048 x 2048 pixels, and the lateral resolution was approximately 1μm without aberration. Since the illumination light sheet was oriented at 45° with respect to the sample surface, the 1mm x 1mm FOV corresponded to 0.67 mm x 1mm in the depth and width axes. A custom LabView program was developed to control the imaging process. The program generated an analog voltage output signal to control the ETL driver, and a digital trigger output signal to initiate the rolling shutter of the sCMOS camera, and a digital output signal to move the microscope stage.

**Fig. 1.**
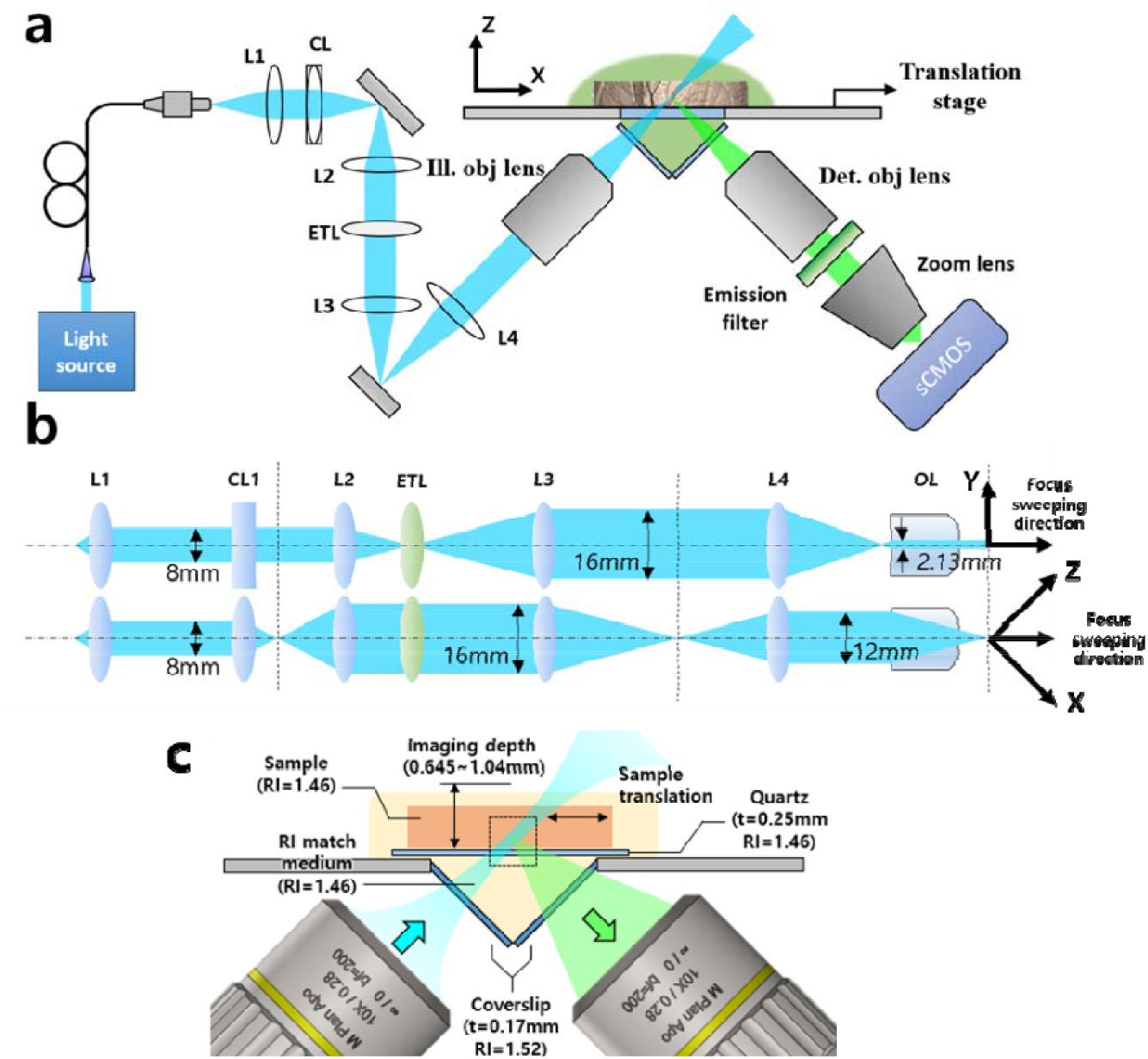
Schematics of OTAS-LSM. (a) an overall schematic of OTAS-LSM, (b) a detail optical configuration in the illumination arm, (c) a detail schematic in the sample and water prism

**Fig. 2.**
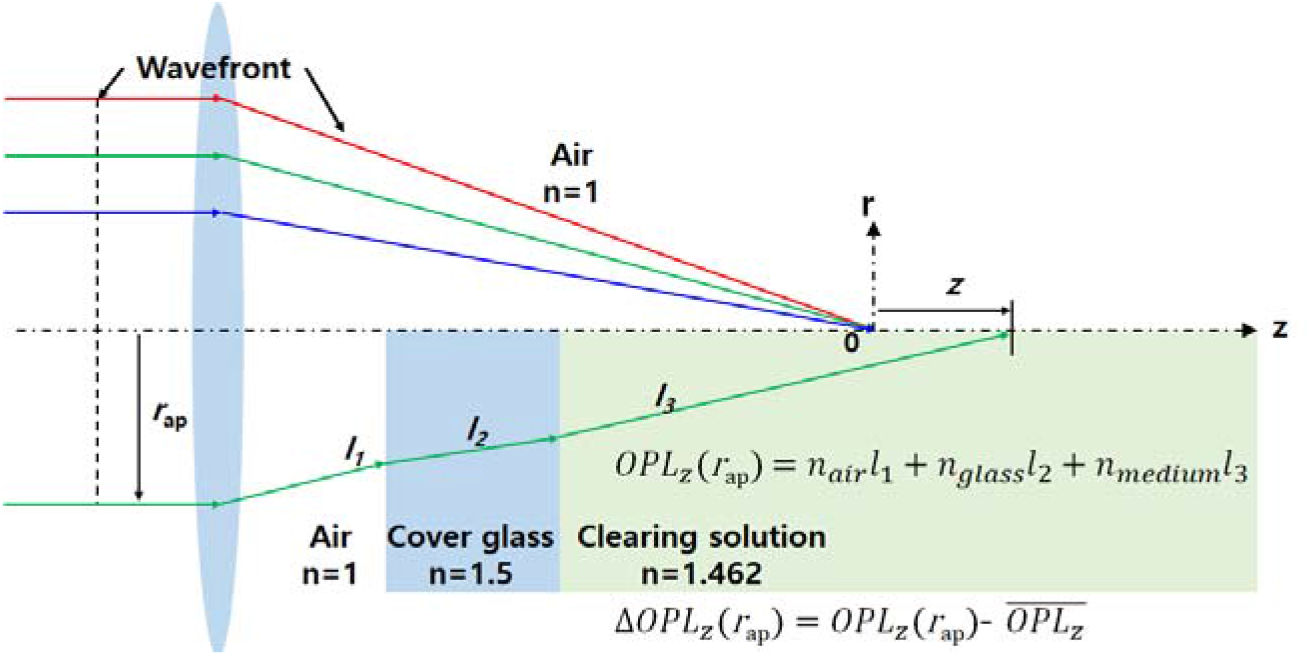
A ray diagram for the aberration analysis of OTAS-LSM. The upper half is the ideal case without RI mismatch, and the lower half is the actual case where light rays travel through the layers of different RIs. A ray launched parallel to the optical axis experienced multiple refractions by going through the layers of different RIs. Path length of the ray was calculated by adding up the individual length segments multiplied with the corresponding RIs.

### 2.2 System characterization with aberration simulation and measurement

The current OTAS-LSM design had on-axis optical aberration by using the air objective lenses and the water prism. The refractive index mismatch of the air and the RI matching solution in the light path caused the aberration. The aberration was analyzed by both simulation and experiment. A custom Matlab code was developed for the simulation. Light rays, which launched in parallel to the optical axis before the back aperture of the objective lens, experienced multiple refractions by passing through the layers of different RIs including the air, coverslip, RI matching solution until arriving at a point (z) in the optical axis. Path length of the ray to the point in the optical axis was calculated by adding individual path length segments multiplied with the corresponding RIs. Path length differences (PLDs) among the rays at different radial positions in the objective back aperture (*r*_ap_) were obtained. Light intensity at the point in the optical axis was calculated by combining the rays. The geometrical coordinate (z=0) was set at the focus of the light rays in the aberration free condition. The axial location of the maximum intensity was found, and it was set to be the focal point in the aberrated condition. The PLDs at the focal point were analyzed by using Zernike polynomials. Dominant Zernike coefficients in the current system were defocus and spherical aberration. The obtained spherical aberration coefficient was used to construct the point spread function (PSF) in the aberrated condition.

The imaging resolution was experimentally measured by imaging fluorescent microspheres (0.5μm in diameter) embedded in the agarose and RI matching solution mixture. The intensity profiles of the microsphere images were obtained and fitted to the Gaussian function for the analysis of full width at half maximum intensity (FWHM). More than 10 microsphere images were analyzed, and average FWHM values were obtained.

### 2.3 Sample preparation of optically cleared tissues and OTAS-LSM imaging

After the characterization, OTAS-LSM was applied to the imaging of optically cleared tissues including the brain slices of Thy1-eYFP mice and the small intestines of ChAT-Cre Tomato TG mice, *ex vivo*. In the preparation of the mouse brain slices, the mouse was euthanized and the whole mouse brain was extracted and fixed in 4% formaldehyde solution for 24 hours. The fixed mouse brain was sliced coronally in 1mm thickness. The brain slices were incubated in the RI matching solution at 36□ for 24 hours for the optical clearing. The RI matching solution increased the tissue transparency by matching the RI of the fixed brain tissue. In case of the small intestine, the excised tissues were incubated in formaldehyde solution for 24 hours and then in the RI matching solution at 36□ for 6 hours.

The optically cleared tissue specimens were mounted in the sample holder of OTAS-LSM and imaged in 3D with the sample translation in both the x and y directions. The tissue specimens were imaged in the unit of image strips which were generated by the translation in the x direction in the step size of 1μm. The size of each image strip was 930μm × 0.63μm in the width and height, and the length varied depending on the sample size. In case of the mouse brain slice, 12 image strips in total were acquired to cover the entire area. Total imaging time was approximately 1 hour (67 minutes). The reconstructed volumetric image was 8.1mm x 11.4mm x 0.67mm in size. In case of the mouse small intestine specimens, 8 image strips in total were acquired. The imaging time was approximately 30 minutes. The reconstructed volumetric image was approximately 4.5mm x 8mm x 0.67mm. The acquired volumetric images of the optically cleared tissues were processed and visualized by using Matlab. Lucy-Richardson deconvolution algorithm was applied by using simulated PSF information.

### 2.4 Confocal microscopy imaging for comparison with OTAS-LSM

The performance of OTAS-LSM was characterized in comparison with confocal microscopy in the mouse brain. Confocal imaging was conducted by using a commercial system (SP-5, Leica). A multi-immersion 20x objective lens (HC PL Apo 20x/0.7 Imm, Leica) was used for the imaging in the RI matching solution. The theoretical image resolution was 0.27μm and 1.83μm in the transverse and axial directions, respectively. The imaging field of view (FOV) was 258μm x 258μm consisting of 1024 x 1024 pixels. Volumetric images were acquired with the axial step size of 1μm and the imaging speed of 2 frames/s. The imaging time for 1mm^3^ volume was approximately 1hr.

## 3. Result

### 3.1 Aberration Analysis of OTAS-LSM

The results of image resolution analysis in OTAS-LSM are shown in Fig. 3. A schematic showing the illumination and imaging light paths and the geometrical coordinate are shown in Fig. 3(a). The simulation and experiment results are shown in Fig. 3(b) and 3(c), respectively. In the simulation, the changes of excitation light sheet, imaging PSF, and system PSF by the aberration were analyzed. The excitation light sheet, which was 0.73μm thick in FWHM in the aberration free condition, was degraded to 1.00μm with the aberration. The imaging PSF, which was 0.79μm in FWHM in the aberration free condition, was degraded to 1.06μm with the aberration. The system PSF could be obtained by combining the axially swept illumination light sheet and imaging PSF which crossed each other just like the ones in the schematic, and the results in both the aberration free and aberrated conditions are shown in Fig. 3(b). The system resolutions, which were 1.14μm in the x-z plane and 0.79μm in the y axis in the aberration free condition, were degraded to 1.50μm, 1.06μm with the aberration, respectively. The image resolution of the OTAS-LSM was degraded to approximately 131% of the theoretical values.

**Fig. 3.**
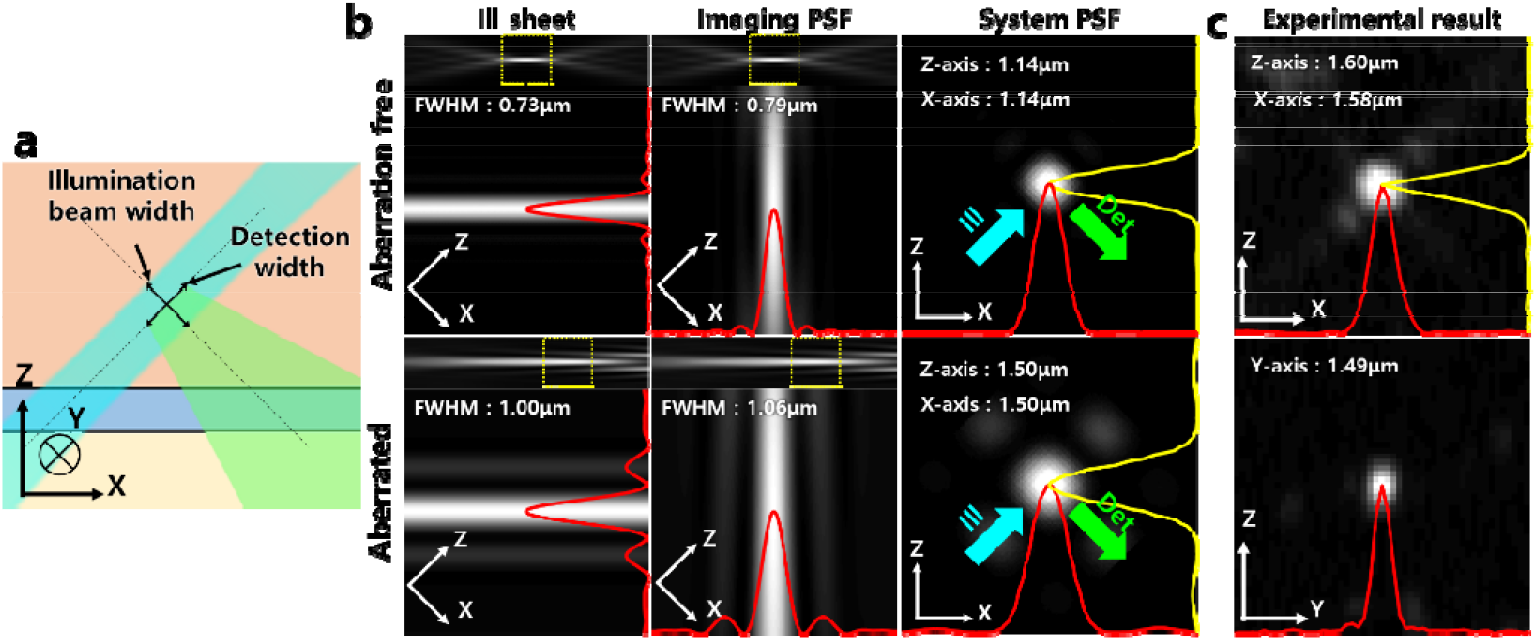
Resolution analysis in both simulation and experiment. (a) a schematic showing the excitation light sheet and emission light paths and the geometrical coordinates (b) Simulation results of excitation light sheet, detection point spread function (PSF), and system PSF in both the aberration-free and aberration conditions. (c) Experiment results showing representative images of a 0.5μm fluorescent microsphere and intensity profiles.

In the experiment, the PSF of microspheres was measured to be 1.60 ± 0.17 μm and 1.58 ± 0.15 μm in FWHM in the x-z plane and 1.49 ± 0.04 μm in the y axis, respectively. The measured image resolutions were 106.6% (x), 105.3% (z) and 140.5% (y) of the simulation results with the aberration, and approximately 140% (x, z) and 188.6% (y) of the ideal theoretical results. The analysis concluded that the image resolutions of OTAS-LSM system were in between 1 to 2μm in both the simulation and experiment.

### 3.2 OTAS-LMS imaging of optically cleared mouse brain and mouse small intestine

After the system characterization, OTAS-LSM was applied to optically cleared tissues including the mouse brain and small intestine for the cell-level visualization of neuronal network. Thy1-eYFP mice expressing eYFP in motor and sensory neurons were used for the imaging of neural network in the brain. ChAT-Cre-tdTomato TG mice expressing tdTomato fluorescent protein in the cholinergic innervation and nonneuronal acetylcholine-synthesizing cells encountered in the periphery were used for the imaging of enteric nerve system in the small intestine [21]. OTAS-LSM images of the mouse brain slice are shown in Fig. 4. An image of the entire brain slice and magnified images in the several regions of interest (ROIs) are shown. These images were volumetric images with depth color coding: the depth range from 0μm to 200μm from the surface for the gross image in Fig. 4(a) and the range from 0μm to 100μm deep for the magnified images in Fig. 4(b-d), respectively. The image of entire mouse brain slice showed gross neural network in the mouse brain: sparse network in the cerebral cortex and dense network in the hippocampus. The magnified images in 3 different ROIs of the mouse brain showed detail neural network in the single cell level. The magnified image in the cerebral cortex (Fig. 4b) showed pyramidal cells with apical dendrites extending upward to the cortical surface. The apical dendrites were branched for several times in the extension. The image in the hippocampus showed pyramidal cells with thin basal dendrites arising from the soma and axons extending downward. The magnified image in the thalamus showed nerve fibers projecting out to the cerebral cortex. Individual nerve fibers were resolved. OTAS-LSM could visualize the neural network of the mouse brain at the resolution of individual neurons and their compartments.

**Fig. 4.**
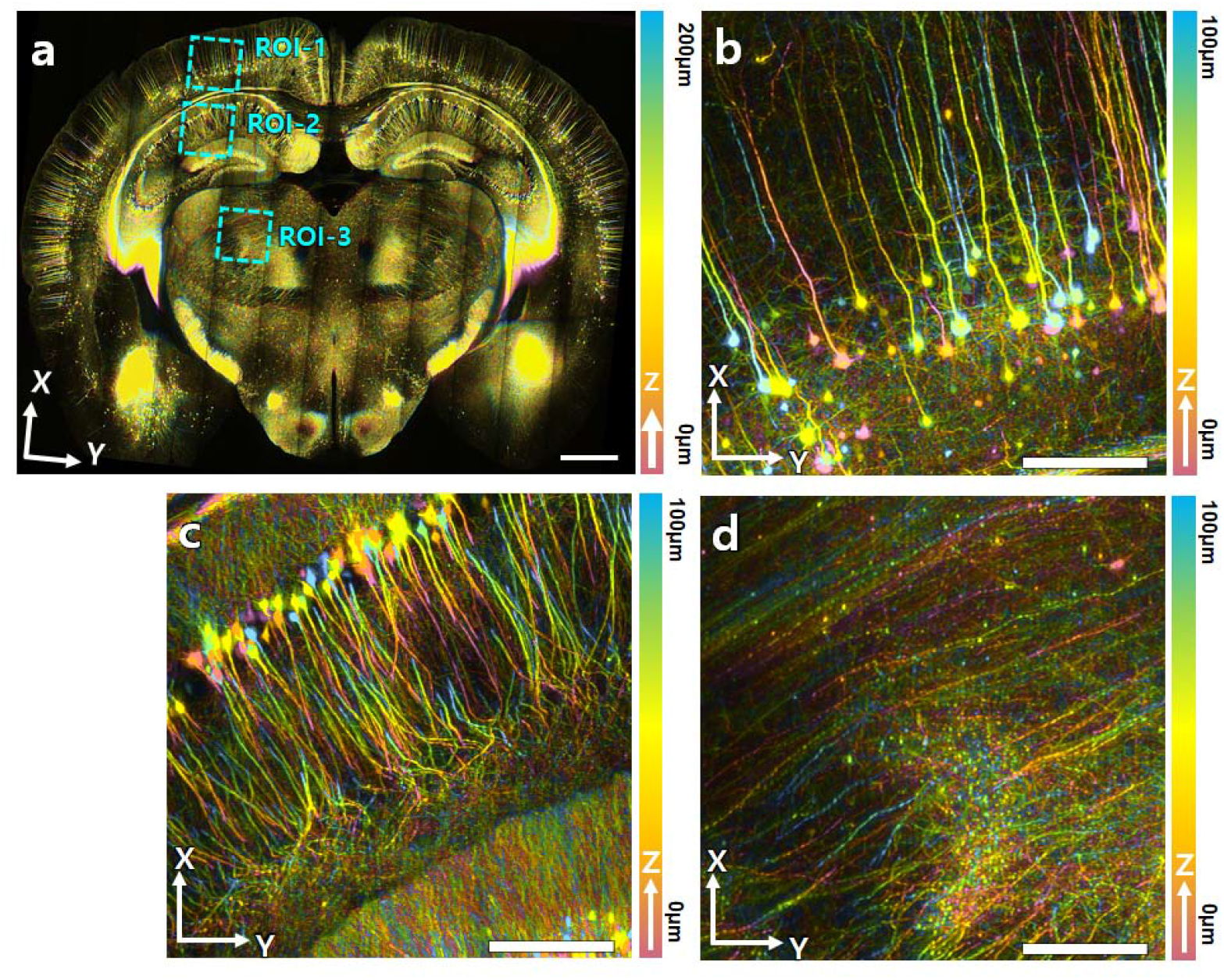
OTAS-LSM images of an optically cleared Thy1-eYFP mouse brain. (a) a large sectional image of the brain slice with depth color coding from 0 to 200μm. (b-d) magnified images of the cerebral cortex, hippocampus, and thalamus with depth color coding from 0 to 60μm, respectively. Scale bars: 1mm in (a), 200μm in (b-d)

OTAS-LSM images of the enteric nerve system (ENS) in the mouse small intestine are shown in Fig. 5. A 3D image in the large FOV of 7mm x 3mm and three magnified images in ROIs are shown. The large FOV image was presented in depth color coding, and the magnified images were presented as the maximum intensity projection (MIP) of the 3D image in specific depth ranges. The large FOV image showed the gross structure of ENS. Neuronal networks in different morphologies or structures appeared at different depth ranges. The magnified images showed the myenteric nerve plexus (0 - 20μm), deep muscular plexus. (20 - 30μm), and submucosal plexus (30μm and more).

**Fig. 5.**
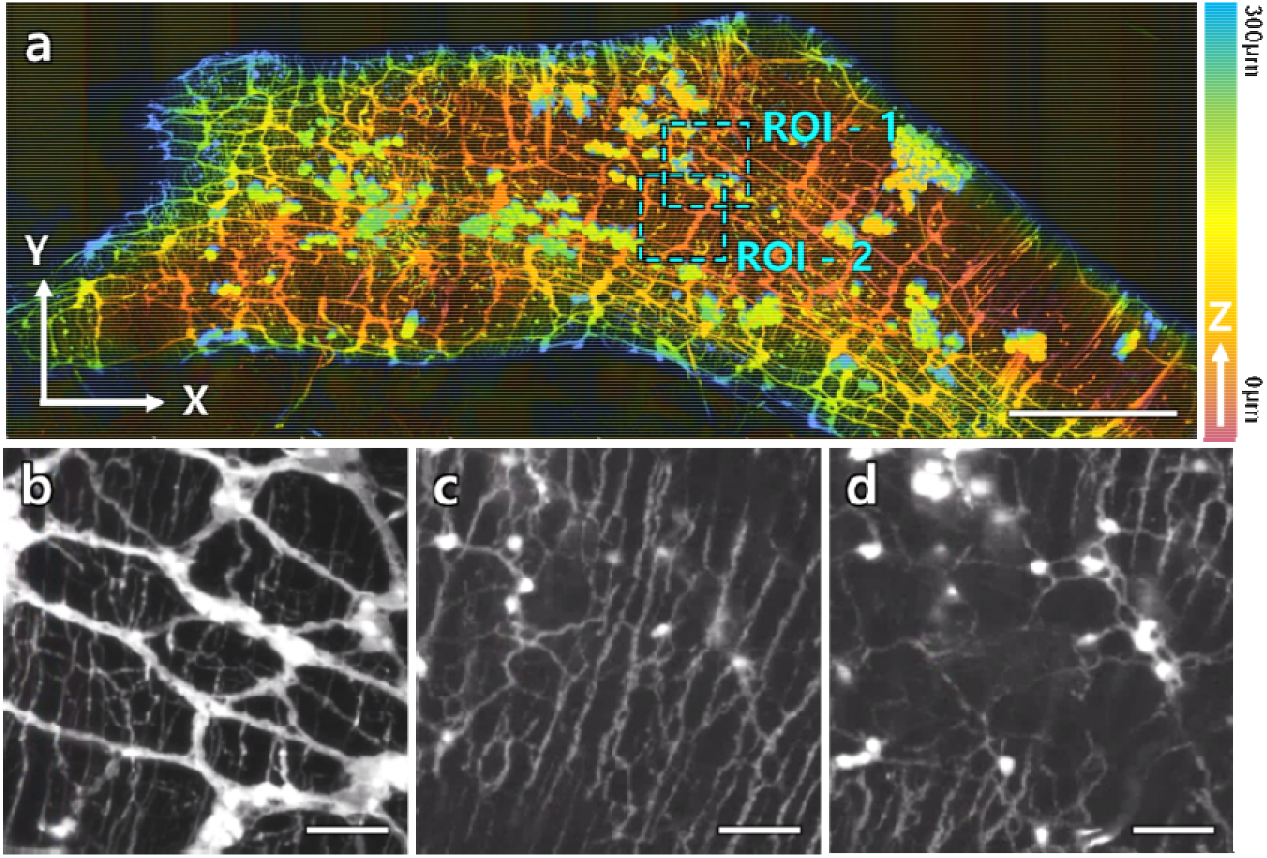
OTAS-LSM images of an optical cleared small intestine of the ChAT-Cre-tdTomato transgenic mouse. (a) a large sectional image with depth color coding from 0 to 300μm. (b-d) magnified images of the region of interests (ROIs) marked in the large sectional image and processed as maximum intensity projection (MIP). (b) an MIP image of ROI-1 showing the myenteric nerve plexus from 0 to 20μm from the surface. (c) and (d) are MIP images of the deep muscular plexus from 20 to 30 μm in depth and submucosal plexus from 30 to 40 μm in depth. Scale bars are 1mm in (a) and 100μm in (b-d).

### 3.3 Comparison of OTAS-LSM with confocal microscopy in the mouse brain

OTAS-LSM was compared with confocal microscopy (CM) in the imaging of mouse brain, and the results are in Fig. 6. Both the OTAS-LSM and CM images in two different regions of the mouse brain, the cerebral cortex (Fig. 6a and b) and hippocampus (Fig. 6c and d), are shown. Both OTAS-LSM and CM images were in the same FOV of 260μm x 260μm. CM visualized fine cellular structures down to thin dendrites (Fig. 6 b, d) with the higher image resolution than those of OTAS-LSM. However, OTAS-LSM could resolved individual neurons at the modest image resolution of 1-2μm (Fig. 6 a, c). The imaging throughputs of CM and OTAS-LSM would be quite different, owing to the difference in the imaging resolution. Additionally, CM needed extra time for switching the imaging sections: approximately 1s per switching. OTAS-LSM could do the 3D imaging continuously via the linear translation of samples without much additional time. Therefore, OTAS-LSM was advantageous for the high throughput visualization of gross cellular structure of the optically cleared tissues.

**Fig. 6.**
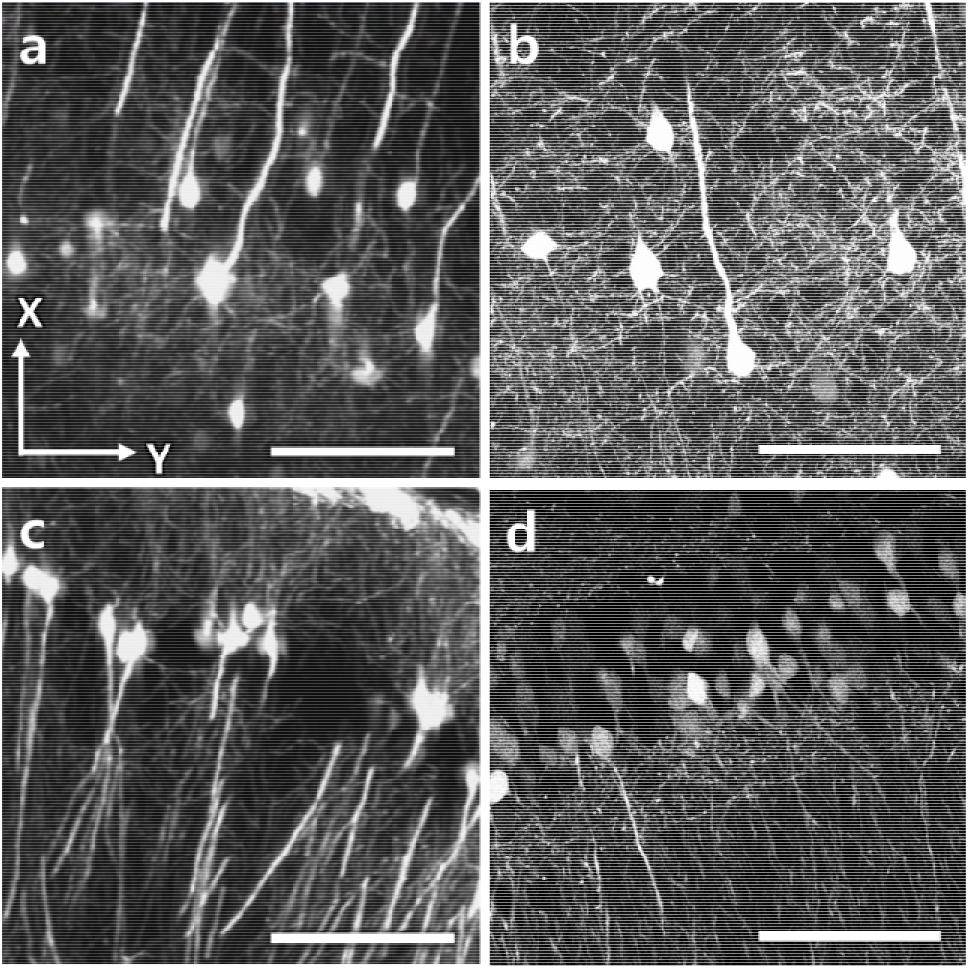
Comparison between OTAS-LSM and confocal microscopy (CM) in the visualization of neuronal network in the brain. (a, b) MIP OTAS-LSM and CM images of the cerebral cortex, respectively. (c, d) MIP OTAS-LSM and CM images of the hippocampus. Scale bars: 100μm.

## 4. Discussion

OTAS-LSM was developed for the high throughput imaging of optically cleared tissues at the improved axial resolution. The current OTAS-LSM was based on the water-prism OT-LSM design for simplicity. OTAS-LSM implemented the axial sweeping of excitation light sheet by using the ETL. The emission light generated at the focus of light sheet was collected confocally by synchronizing the rolling shutter in the imaging camera with the excitation light sheet. The imaging FOV was approximately 1mm x 1mm in the oblique imaging plane, and the imaging speed was 30 frames/s. Although the current OTAS-LSM had the inherent on-axis optical aberration by using the air objective lenses and the water prism, the image resolution was within 1-2μm in both the axial and transverse directions. In the imaging of optically cleared mouse brain and small intestine tissues, OTAS-LSM demonstrated its ability of visualizing the neural network in the mouse brain and the enteric nerve system in the small intestine in the single cell level.

The current OTAS-LSM had several limitations including the imaging depth range, the image resolution, and the imaging speed. The current system could image the sample down to less than 1 mm deep from the surface, because the sample holder translated the sample in the x-y directions only. A new sample holder, which could axially translate the sample, would increase the imaging depth range [<u>ref</u>]. The new sample holder is currently under development. The current system had the on-axis optical aberration, and the aberration would be worse with the increase of the imaging depth. Aberration correction methods need to be implemented. Curved immersion meniscus lenses could replace the current water prism with flat coverslips [17]. Wavefront correction devices such as a deformable mirror (DM) or a spatial light modulator (SLM) could be incorporated [14, 22, 23]. The current imaging speed was up to 30 frames/s limited by the ETL. Other devices such as deformable mirrors, which have the faster response, could be used both to axially sweep the light sheet and to correct aberration.

## 5. Conclusion

OTAS-LSM was developed for the high-throughput cellular imaging of optically cleared large tissues at the improved axial resolution. OTAS-LSM was implemented by using an ETL for the axial sweeping, and air objective lenses and the water prism for illumination and imaging. The image resolution of OTAS-LSM was 1 to 2 μm with some degradation associated with the on-axis optical aberration. The performance of OTAS-LSM was verified in the imaging of the optically cleared mouse brain and small intestine and it could visualize neuronal network in the cell level. OTAS-LSM might be useful for the cellular examination of optically cleared tissues by providing cellular information at high throughputs.

## References

1. J. Huisken, J. Swoger, F. Del Bene, J. Wittbrodt, and E. H. K. Stelzer, “Optical sectioning deep inside live embryos by selective plane illumination microscopy,” Science 305, 1007–1009 (2004).

2. P. J. Keller, A. D. Schmidt, J. Wittbrodt, and E. H. Stelzer, “Reconstruction of zebrafish early embryonic development by scanned light sheet microscopy,” Science 322, 1065–1069 (2008).

3. M. B. Ahrens, M. B. Orger, D. N. Robson, J. M. Li, and P. J. Keller, “Whole-brain functional imaging at cellular resolution using light-sheet microscopy,” Nature Methods 10, 413–+ (2013).

4. H. U. Dodt, U. Leischner, A. Schierloh, N. Jahrling, C. P. Mauch, K. Deininger, J. M. Deussing, M. Eder, W. Zieglgansberger, and K. Becker, “Ultramicroscopy: three-dimensional visualization of neuronal networks in the whole mouse brain,” Nat Methods 4, 331–336 (2007).

5. K. M. Dean, P. Roudot, E. S. Welf, G. Danuser, and R. Fiolka, “Deconvolution-free Subcellular Imaging with Axially Swept Light Sheet Microscopy,” Biophys J 108, 2807–2815 (2015).

6. P. N. Hedde and E. Gratton, “Selective plane illumination microscopy with a light sheet of uniform thickness formed by an electrically tunable lens,” Microsc Res Tech 81, 924–928 (2018).

7. T. Chakraborty, M. K. Driscoll, E. Jeffery, M. M. Murphy, P. Roudot, B. J. Chang, S. Vora, W. M. Wong, C. D. Nielson, H. Zhang, V. Zhemkov, C. Hiremath, E. D. De La Cruz, Y. Yi, I. Bezprozvanny, H. Zhao, R. Tomer, R. Heintzmann, J. P. Meeks, D. K. Marciano, S. J. Morrison, G. Danuser, K. M. Dean, and R. Fiolka, “Light-sheet microscopy of cleared tissues with isotropic, subcellular resolution,” Nat Methods 16, 1109–1113 (2019).

8. J. Ping, F. Zhao, J. Nie, T. Yu, D. Zhu, M. Liu, and P. Fei, “Propagating-path uniformly scanned light sheet excitation microscopy for isotropic volumetric imaging of large specimens,” J Biomed Opt 24, 1–5 (2019).

9. F. F. Voigt, D. Kirschenbaum, E. Platonova, S. Pages, R. A. A. Campbell, R. Kastli, M. Schaettin, L. Egolf, A. van der Bourg, P. Bethge, K. Haenraets, N. Frezel, T. Topilko, P. Perin, D. Hillier, S. Hildebrand, A. Schueth, A. Roebroeck, B. Roska, E. T. Stoeckli, R. Pizzala, N. Renier, H. U. Zeilhofer, T. Karayannis, U. Ziegler, L. Batti, A. Holtmaat, C. Luscher, A. Aguzzi, and F. Helmchen, “The mesoSPIM initiative: open-source light-sheet microscopes for imaging cleared tissue,” Nat Methods 16, 1105–1108 (2019).

10. B. Migliori, M. S. Datta, C. Dupre, M. C. Apak, S. Asano, R. Gao, E. S. Boyden, O. Hermanson, R. Yuste, and R. Tomer, “Light sheet theta microscopy for rapid high-resolution imaging of large biological samples,” BMC Biol 16, 57 (2018).

11. V. Voleti, K. B. Patel, W. Li, C. Perez Campos, S. Bharadwaj, H. Yu, C. Ford, M. J. Casper, R. W. Yan, W. Liang, C. Wen, K. D. Kimura, K. L. Targoff, and E. M. C. Hillman, “Real-time volumetric microscopy of in vivo dynamics and large-scale samples with SCAPE 2.0,” Nat Methods 16, 1054–1062 (2019).

12. M. B. Bouchard, V. Voleti, C. S. Mendes, C. Lacefield, W. B. Grueber, R. S. Mann, R. M. Bruno, and E. M. Hillman, “Swept confocally-aligned planar excitation (SCAPE) microscopy for high speed volumetric imaging of behaving organisms,” Nat Photonics 9, 113–119 (2015).

13. R. McGorty, H. Liu, D. Kamiyama, Z. Dong, S. Guo, and B. Huang, “Open-top selective plane illumination microscope for conventionally mounted specimens,” Opt Express 23, 16142–16153 (2015).

14. R. McGorty, D. Xie, and B. Huang, “High-NA open-top selective-plane illumination microscopy for biological imaging,” Opt Express 25, 17798–17810 (2017).

15. A. K. Glaser, N. P. Reder, Y. Chen, E. F. McCarty, C. Yin, L. Wei, Y. Wang, L. D. True, and J. T. C. Liu, “Light-sheet microscopy for slide-free non-destructive pathology of large clinical specimens,” Nat Biomed Eng 1(2017).

16. Y. Chen, W. Xie, A. K. Glaser, N. P. Reder, C. Mao, S. M. Dintzis, J. C. Vaughan, and J. T. C. Liu, “Rapid pathology of lumpectomy margins with open-top light-sheet (OTLS) microscopy,” Biomed Opt Express 10, 1257–1272 (2019).

17. L. A. Barner, A. K. Glaser, L. D. True, N. P. Reder, and J. T. C. Liu, “Solid immersion meniscus lens (SIMlens) for open-top light-sheet microscopy,” Opt Lett 44, 4451–4454 (2019).

18. L. A. Barner, A. K. Glaser, H. Huang, L. D. True, and J. T. C. Liu, “Multi-resolution open-top light-sheet microscopy to enable efficient 3D pathology workflows,” Biomed Opt Express 11, 6605–6619 (2020).

19. A. K. Glaser, N. P. Reder, Y. Chen, C. Yin, L. Wei, S. Kang, L. A. Barner, W. Xie, E. F. McCarty, C. Mao, A. R. Halpern, C. R. Stoltzfus, J. S. Daniels, M. Y. Gerner, P. R. Nicovich, J. C. Vaughan, L. D. True, and J. T. C. Liu, “Multi-immersion open-top light-sheet microscope for high-throughput imaging of cleared tissues,” Nat Commun 10, 2781 (2019).

20. A. K. Glaser, K. W. Bishop, L. A. Barner, R. B. Serafin, and J. T. C. Liu, “A hybrid open-top light-sheet microscope for multi-scale imaging of cleared tissues,” BioRxiv (2020).

21. L. Gautron, J. M. Rutkowski, M. D. Burton, W. Wei, Y. Wan, and J. K. Elmquist, “Neuronal and nonneuronal cholinergic structures in the mouse gastrointestinal tract and spleen,” J Comp Neurol 521, 3741–3767 (2013).

22. C. Bourgenot, C. D. Saunter, J. M. Taylor, J. M. Girkin, and G. D. Love, “3D adaptive optics in a light sheet microscope,” Optics Express 20, 13252–13261 (2012).

23. D. Wilding, P. Pozzi, O. Soloviev, G. Vdovin, and M. Verhaegen, “Adaptive illumination based on direct wavefront sensing in a light-sheet fluorescence microscope,” Optics Express 24, 24896–24906 (2016).

